# Housing temperature dictates the systemic and tissue-specific molecular responses to cancer in mice

**DOI:** 10.1101/2024.05.29.596034

**Authors:** Andrea Irazoki, Emma Frank, Tang Cam Phung Pham, Jessica L Braun, Amy M. Ehrlich, Mark Haid, Fabien Riols, Camilla Hartmann Friis Hansen, Anne-Sofie Rydal Jørgensen, Nicoline Resen Andersen, Laura Hidalgo-Corbacho, Roberto Meneses-Valdes, Mona Sadek Ali, Steffen Henning Raun, Johanne Louise Modvig, Samantha Gallero, Steen Larsen, Zach Gerhart-Hines, Thomas Elbenhardt Jensen, Maria Rohm, Jonas T. Treebak, Val Andrew Fajardo, Lykke Sylow

**Affiliations:** Department of Biomedical Sciences, Faculty of Health and Medical Sciences, University of Copenhagen, Denmark; Department of Kinesiology, Faculty of Applied Health Sciences, Cairns Family Health and Bioscience Research Complex, Brock University, Niagara Region, Ontario, Canada; Novo Nordisk Foundation Center for Basic Metabolic Research, Faculty of Health and Medical Sciences, University of Copenhagen, Denmark; Metabolism & Proteomics Core, Helmholtz Center Munich, German Research Center for Environmental Health, Neuherberg, Germany; Section of Experimental Animal Models, Department of Veterinary and Animal Sciences, University of Copenhagen, Denmark; Department of Nutrition, Exercise and Sport, University of Copenhagen, Denmark; Clinical Research Centre, Medical University of Bialystok, Poland; Institute of Sports Medicine Copenhagen, Department of Orthopedic Surgery M, Copenhagen University Hospital – Bispebjerg and Frederiksberg, Denmark; Institute for Diabetes and Cancer, Helmholtz Center Munich, Neuherberg, Germany; Joint Heidelberg-IDC Translational Diabetes Program, Inner Medicine 1, Heidelberg University Hospital, Heidelberg, Germany; German Center for Diabetes Research (DZD), Neuherberg, Germany

**Keywords:** cancer cachexia, cold-induced stress, thermoneutrality, thermogenic tissues, bioenergetics

## Abstract

Cancer cachexia is a metabolic condition affecting up to 80% of patients with cancer. Cachexia is mediated by reduced muscle and fat mass and impaired function, and it lowers survival for patients. With no approved drugs to treat cachexia, preclinical efforts focus on understanding the molecular mechanisms underlying this condition to reveal treatment targets. Housing laboratory mice at ambient temperature imposes cold stress, leading to induced thermogenic activity and consequent whole-body metabolic adaptations. Yet, the impact of housing temperature in *in vivo* preclinical cachexia remains unknown. We found that thermoneutral (TN) housing in C26 carcinoma-bearing (C26) mice affected lean and fat mass, but not muscle weight or force. TN housing improved glucose tolerance in C26 mice, while enhancing circulating abundance of FGF21 and IL-6. Thermogenic tissues, especially brown adipose tissue, exhibited housing temperature-dependent molecular responses to cancer in oxygen consumption, ATP levels and SERCA ATPase activity, which are all crucial for cancer-induced whole-body metabolic adaptations. We conclude that molecular and systemic adaptations to cancer in mice critically depend on housing temperature, which should be considered in the design and interpretation of preclinical cancer studies.

## Main

Cancer cachexia (CC) is a metabolic condition affecting up to 80% of patients with cancer^1^, mediated by reduced skeletal muscle (SkM) and fat mass and impaired function^2^. It dramatically compromises quality of life, increases treatment toxicity and lowers survival rate. Despite exciting recent discoveries obtained using mouse models that describe potentially targetable molecular mechanisms, the enthusiasm is tempered by their failure to fully replicate the human condition^3^. Unlike humans, whose thermoneutral zone lies at ambient temperature, housing laboratory mice at ambient or standard temperature (ST; 20-22°C) results in metabolic adaptations driven by cold stress-induced thermogenic activity, which do not occur when housing mice at their thermoneutral zone (TN; 28-32°C^4^). In fact, we and others have shown that, compared to TN housing, ST housing affects metabolic adaptations in preclinical models of obesity^5^, insulin action^6^, and in response to exercise^7,8^, showing that housing mice under cold stress represents a substantial confounding factor. Albeit CC is considered a metabolic condition, the impact of housing temperature on murine CC remains unexplored. Thus, in this study we assessed the contribution of cold-induced thermogenic activity to the overall murine CC phenotype by housing tumor-bearing mice at ST or TN conditions and analyzing the whole-body and tissue-specific molecular responses to cancer.

Eighteen to twenty-one days after subcutaneous inoculation of C26-colon carcinoma cells, we observed 8% body weight loss in all C26 mice irrespective of housing temperature (Fig. 1a), changes in food intake (Sup. Fig. 1a) and tumor weight (Sup Fig. 1b). This corresponds to moderate cachexia as previously defined as body weight loss up to 10%^9^. CC is characterized by reduced SkM mass and function. In this regard, only C26 mice housed at TN exhibited decreased lean mass (Fig. 1b), although SkM weight was preserved in all conditions (Sup. Fig. 1c). Despite no significant reduction in grip strength (Sup. Fig. 1d), *ex vivo* force assessment showed substantially decreased force in C26 mice independently of temperature (−42%; Fig. 1c). We also found upregulated atrophy-related gene (atrogenes) expression (Fig. 1d), indicating that muscle weakness and atrophy precede loss of SkM mass upon cancer irrespective of housing temperature. CC is also characterized by alterations in fat mass and function^10^. Notably, only C26 mice housed at TN showed a reduction in their total fat mass (Fig. 1e) compared to their controls, explained by an average 36% decrease gonadal and subcutaneous white adipose tissue weight (gWAT and scWAT, respectively), whereas C26 mice at ST only showed decrease compared to their controls (−12%, Sup. Fig. 1e). The interscapular brown adipose tissue (iBAT, hereafter BAT) showed the same trend (Sup. Fig. 1f). As a readout for adipose tissue function, we evaluated mRNA levels of key genes involved in WAT and BAT homeostasis. We analyzed *Cebpa, Cebpd* (markers for late and early stage adipogenesis, respectively^11^) and *Leptin* (marker of fat storage^12^) in WAT. In BAT, we analyzed *Dio2, Elvol3* and *Sirt7*, essential in driving cold-induced thermogenesis^13^. We observed that cancer, and not temperature, promoted an mRNA expression profile associated with reduced adipogenic capacity and fat storage in WAT (Fig. 1f). In contrast, temperature, and not cancer, markedly reduced the expression of thermogenesis-related genes in BAT (Fig. 1f). To further assess the thermogenic capacity in these conditions, we assessed the expression levels of Uncoupling protein 1 (UCP1), the master regulator of nonshivering thermogenesis in adipose tissues^14^. UCP1 uncouples ATP production from oxidative phosphorylation, a process that dissipates heat. Aligned with previous studies^15^, C26 mice housed at ST displayed a 27%, 88% and 45% downregulation of UCP1 mRNA levels in scWAT, gWAT and BAT, respectively. Interestingly, these reductions were more enhanced at TN (Fig. 1g). Concomitantly, C26 mice at ST showed no changes in BAT UCP1 protein content, whereas TN housing promoted a reduction (−67%, Sup. Fig. 1g). Thus, especially at TN, cancer resulted in whitening of adipose tissue, altering its thermogenic capacity, which contrasts with previous studies undertaken at ST showing induction of browning in CC^10,16^. Altogether, our data indicate that housing temperature differently influences cancer-dependent adaptations especially in BAT of C26 mice with moderate cachexia, which potentially impacts the overall metabolic status.

**Figure 1:**
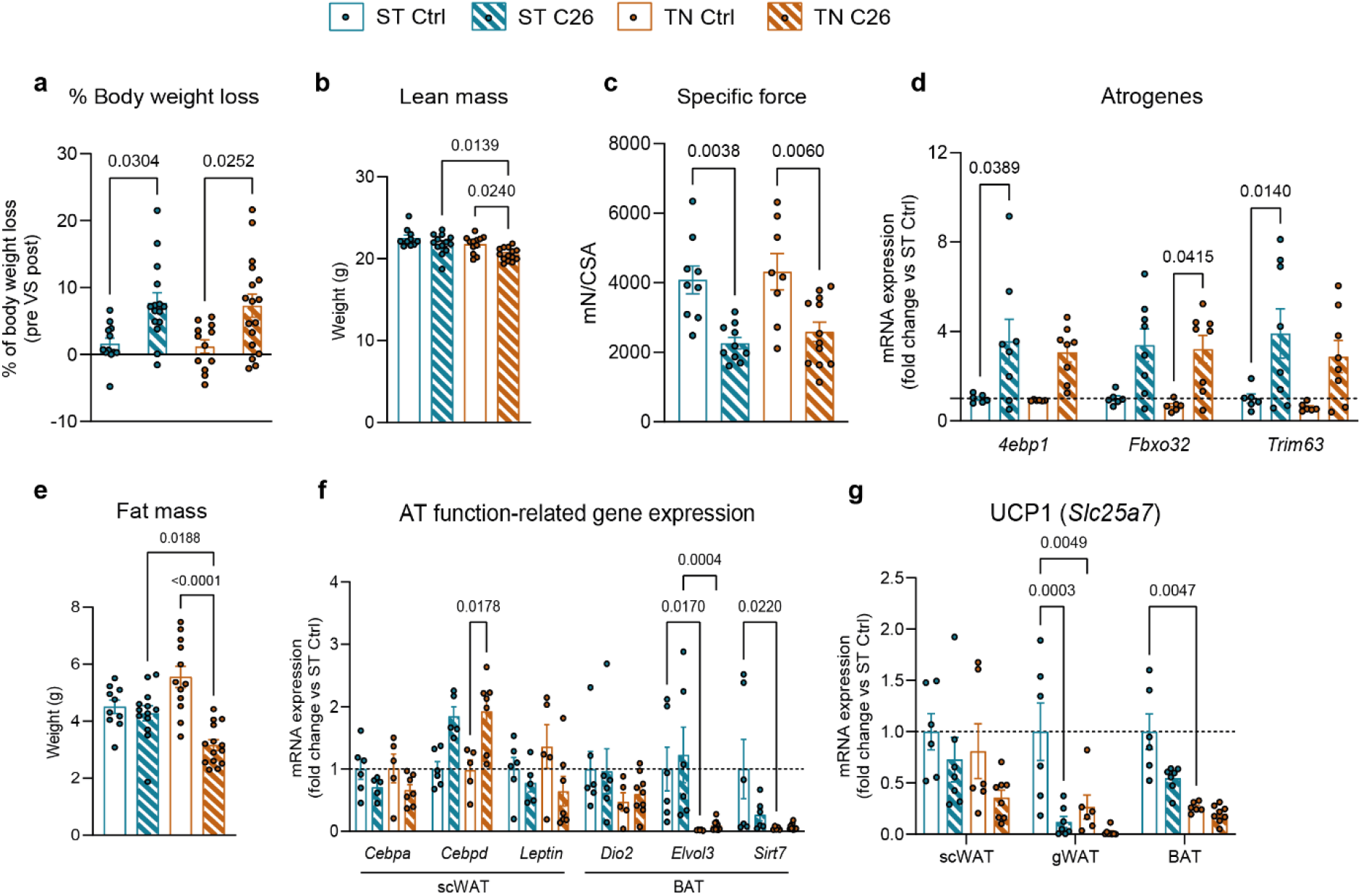
Housing temperature is a critical determinant of cancer’s effect on body composition and fat function, but not muscle force or atrophy. (**a**) Percentage of tumor-free body weight loss comparing the body weight right before C26 cell inoculation (pre) and right before dissection (post). (**b**) Tumor-free lean mass. (**c**) Specific force in soleus muscles. (**d**) Atrogene mRNA expression. (**e**) Fat mass. (**f**) WAT and BAT-related gene expression. (**g**) UCP1 gene expression. Data are expressed as mean ± SE including individual values where applicable. Significance level was set at α=0.05 and only *p*<0.05 are shown. Main effects and interactions were calculated using two-way ANOVA and are shown only when Tukey’s post-hoc test analyses provided no statistical significance between experimental groups. (**a-g**) Two-way ANOVA test with Tukey’s post-hoc test.

Impaired glucose metabolism has been widely reported in response to cancer in mice^17,18^. To assess glycemic control, we subjected the mice to a glucose tolerance test. Aligning with previous studies^17^, C26 mice with moderate cachexia housed at ST displayed glucose intolerance compared to controls (Fig 2a), along with elevated fasting blood glucose and insulin (+17% and +77%, respectively) (Sup. Fig. 2a and b). Strikingly, when housed at TN, C26 mice showed improved glucose tolerance (Fig. 2a), unaltered fasting blood glucose and lowered fasting insulin (Sup. Fig. 2a and b) compared to C26 mice at ST or control mice at TN. These data suggest a potential contribution of cold-induced metabolism in thermogenic tissues to the adaptations in glycemic control upon cancer. Lipid metabolism is also strongly influenced by cancer^10,19^. Therefore, we evaluated temperature-dependent plasma lipid responses to cancer using lipidomics. Principal component analyses (PCA) of plasma showed clustering of cancer-dependent lipid composition (Fig. 2b). Recapitulating previous studies^19^, all C26 mice showed enrichment in circulating modified ceramides, phenocopying human CC (Sup. Fig. 2c). However, the effect of cancer on lipid metabolism seemed independent of housing temperature.

**Figure 2:**
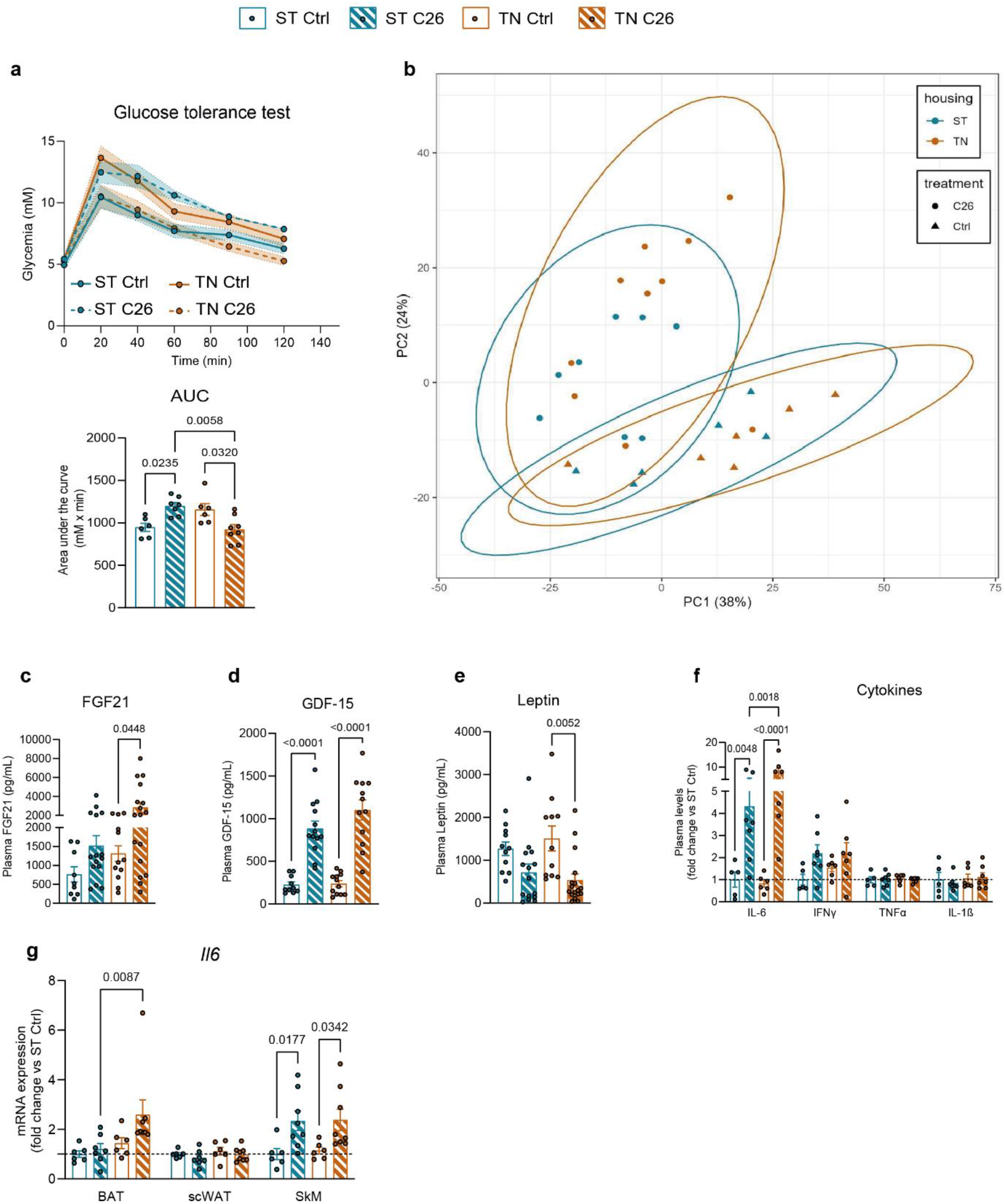
Glucose metabolism and circulating factors, but not lipid composition, are influenced by housing temperature upon cancer. (**a**) Glycemic response during a GTT and average area under the curve (ST control n=6, ST C26 n=7, TN control n=6, TN C26 n=8). (**b**) Principal component analysis (PCA) scores plot of plasma lipid species concentrations. (**c, d, e, f**) Plasma levels of circulating factors. (**g**) IL-6 gene expression. Data are expressed as mean ± SE including individual values where applicable. Significance level was set at α=0.05 and only *p*<0.05 are shown. Main effects and interactions were calculated using two-way ANOVA and are shown only when Tukey’s post-hoc test analyses provided no statistical significance between experimental groups. (**a, c-g**) Two-way ANOVA with Tukey’s post-hoc test.

Both metabolic dysfunction and cancer are characterized by changes in circulating factors, such as fibroblast growth factor 21 (FGF21^20^), growth differentiating factor 15 (GDF-15^21^), leptin^12^ and inflammatory cytokines^22^. C26 mice exhibited an average 4-fold increase in plasma FGF21 (Fig. 2c) and GDF-15 (Fig. 2d) levels compared to their respective control mice, and this effect was augmented for FGF21 at TN by 86% compared to ST. Circulating leptin levels were reduced in all C26 mice independently of housing temperature (Fig. 2e). We also found that C26 mice presented systemic inflammation, indicated by increased circulating IFN-γ and IL-6 (Fig. 2f). Interestingly, circulating IL-6 levels were 76% elevated in C26 mice housed at TN compared to ST. Thermogenic tissues are major sources of IL-6, thus, we hypothesized that changes in housing temperature might affect the cancer-induced IL-6 production in such tissues. Our data demonstrated that SkM, but not adipose tissues, contributed to the IL-6 mediated inflammatory response in C26 mice at ST, whereas at TN, both BAT and SkM produced this cytokine (Fig. 2g). This indicates that housing temperature differentially influences the IL-6 response to cancer from thermogenic tissues, especially in BAT and SkM.

Cellular bioenergetics govern the thermogenic responses to cold stress in thermogenic tissues, and alterations in these processes have the potential to affect the overall metabolic status. Thus, we first evaluated the impact of ST or TN housing on molecular processes involved in cellular bioenergetics in SkM. First, as a functional readout of mitochondrial respiratory capacity, we evaluated O_2_ consumption in permeabilized SkM fibers. In C26 mice housed at ST, we observed a tendency for an increased O_2_ consumption in SkM fibers compared to controls (Fig. 3a). Yet, in mice housed at TN, cancer seemed to lower SkM O_2_ consumption compared to controls. To investigate the outcome of these slight changes in O_2_ consumption, we measured SkM [ATP]. Intriguingly, only C26 mice housed at TN showed a reduction in SkM [ATP] (Fig. 3b). These alterations were independent of changes in mitochondrial mass, measured as TOMM20 protein content (Sup. Fig 3a) and mRNA expression of mitochondrial biogenesis markers *Tfam* and *Ppargc1a* (PGC1α) (Sup. Fig. 3b).

**Figure 3:**
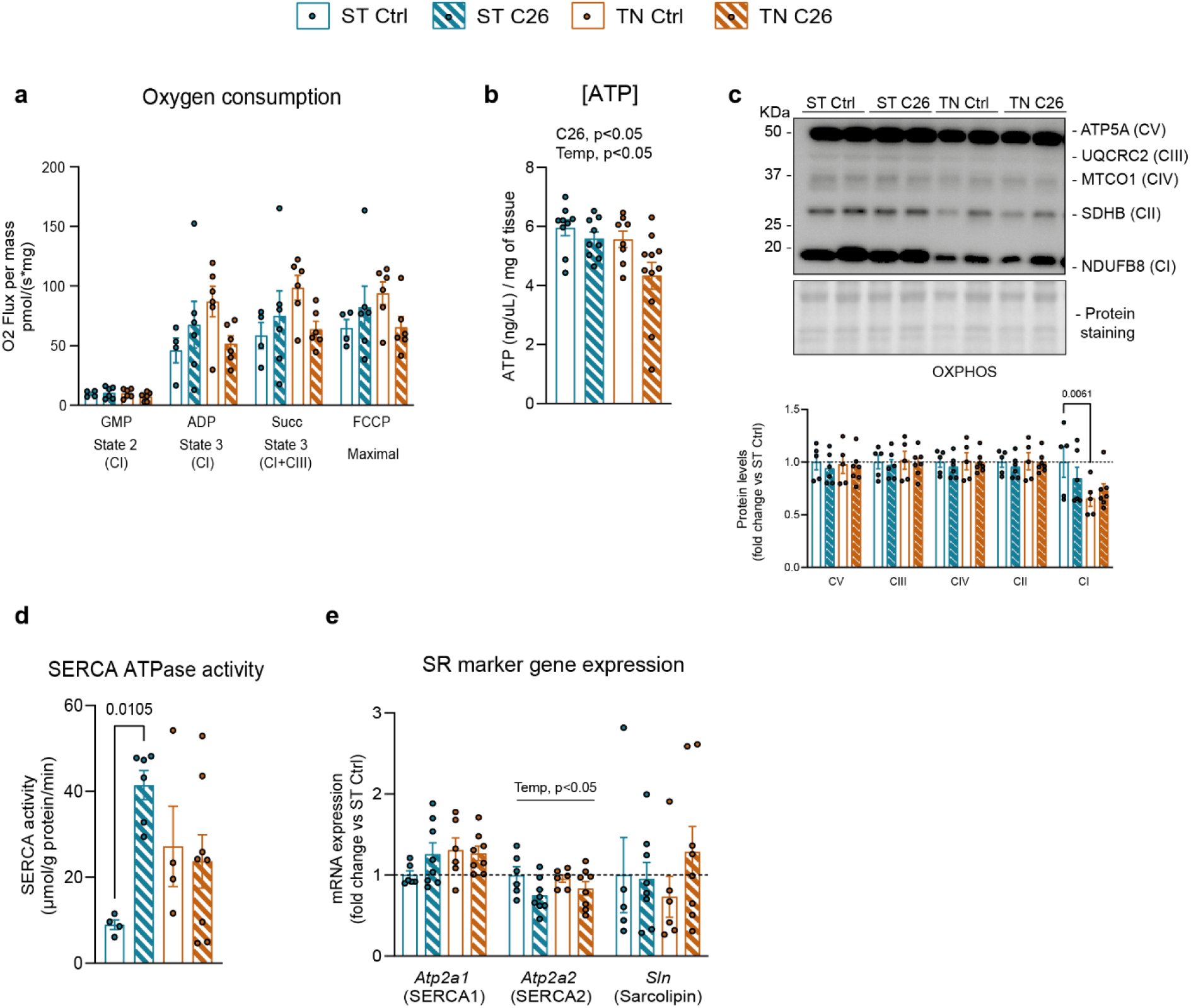
Housing temperature has a mild impact on cancer-induced SkM molecular signature. (**a**) Oxygen consumption in SkM fibers. (**b**) ATP concentration. (**c**) OXPHOS subunit representative immunoblots and quantification. (**d**) SERCA ATPase activity. (**e**) mRNA levels of SR markers. Data are expressed as mean ± SE including individual values where applicable. Significance level was set at α=0.05 and only *p*<0.05 are shown. Main effects and interactions were calculated using two-way ANOVA and are shown only when Tukey’s post-hoc test analyses provided no statistical significance between experimental groups. (**a-e**) Two-way ANOVA test with Tukey’s post-hoc test. C26, main cancer effect; Temp, main temperature effect.

Furthermore, protein quantification of different subunits of the OXPHOS complex showed no changes upon cancer or temperature (Fig 3c), with the only exception of complex I (CI), which was 34% lowered by TN. To further understand the housing temperature-dependent effects on ATP levels upon cancer in SkM, we evaluated the activity of the sarco/endoplasmic reticulum Ca^2+^ ATPase (SERCA ATPase), which pumps Ca^2+^ ions from the cytosol to the sarco/endoplasmic reticulum (SR/ER). This requires ATP hydrolysis, and its uncoupling via sarcolipin results in heat dissipation, a key mechanism in nonshivering thermogenesis in SkM^23^. At ST housing, SkM SERCA ATPase activity was increased 4-fold in C26 mice, while at TN, cancer did not influence SkM SERCA ATPase activity (Fig. 3d). Interestingly, TN housing of control mice seemed to increase SkM SERCA ATPase activity compared to control mice housed at ST. Thus, our data suggest that both cancer and TN housing result in enhanced SERCA ATPase activity in SkM. These changes occurred independently of mRNA levels of SR markers SERCA1, SERCA2 and Sarcolipin (Fig. 3e), and protein levels of SERCA1 and SERCA2 (Sup. Fig. 3c). Notably, we observed a TN-induced reduction in protein levels of STIM1, an SR/ER-residing Ca^2+^ sensor (Sup. Fig. 3c). Reduced STIM1 levels have been associated with decreased mitochondria-free Ca^2+^ and respiration^24^, which suggests that the underlying molecular systems leading to increased SERCA ATPase activity upon cancer or changes in housing temperature are different.

Next, we evaluated the cellular bioenergetics of BAT. We observed a 2-fold increase in O_2_ consumption in permeabilized BAT of C26 mice housed at ST, which was completely blunted at TN (Fig. 4a). Yet, all C26 mice presented increased BAT [ATP] (Fig. 4b), suggesting that the molecular processes involved in ATP synthesis and/or hydrolysis upon cancer depends on housing temperature. These adaptations were not due to alterations in mitochondrial mass, with only a modest increase in TOMM20 protein upon cancer regardless of temperature (+15%, Sup. Fig. 4a), accompanied by a decreasing trend of *Tfam* and *Ppargc1a* (Sup. Fig. 4b). Unlike in SkM, BAT protein levels of subunits of CI, CII and CIII increased a 28% in C26 mice housed at TN, but not at ST (Fig. 4c). These data indicate that bioenergetics of BAT display markedly different cancer-induced phenotypes depending on housing temperature. To evaluate the biology of ATP levels in BAT and UCP1-independent nonshivering thermogenesis, we also assessed SERCA ATPase activity in this tissue. Unlike in SkM, BAT of C26 mice housed at ST showed a 29% reduction in SERCA ATPase activity compared to controls, and this reduction was enhanced at TN (Fig. 4d). Indeed, the exacerbated reduction observed at TN was due to the fact that control mice at TN exhibited a marked increase compared to controls at ST. Thus, TN housing exerts a major impact on BAT SERCA ATPase activity, and this effect is blunted by cancer. Interestingly, SERCA2 mRNA levels tended to be increased in C26 mice housed at ST, but not at TN, when compared to controls (Fig. 4e), although its protein levels in all C26 mice were reduced (−14%, Sup. Fig. 4c). Unlike in SkM, STIM1 was unaltered upon cancer or temperature in BAT. Our data demonstrate that increased BAT ATP levels in C26 mice housed at TN are not due to decreased SERCA-mediated ATP hydrolysis. Therefore, the cancer-induced effects on ATP regulation depended on temperature in BAT.

**Figure 4:**
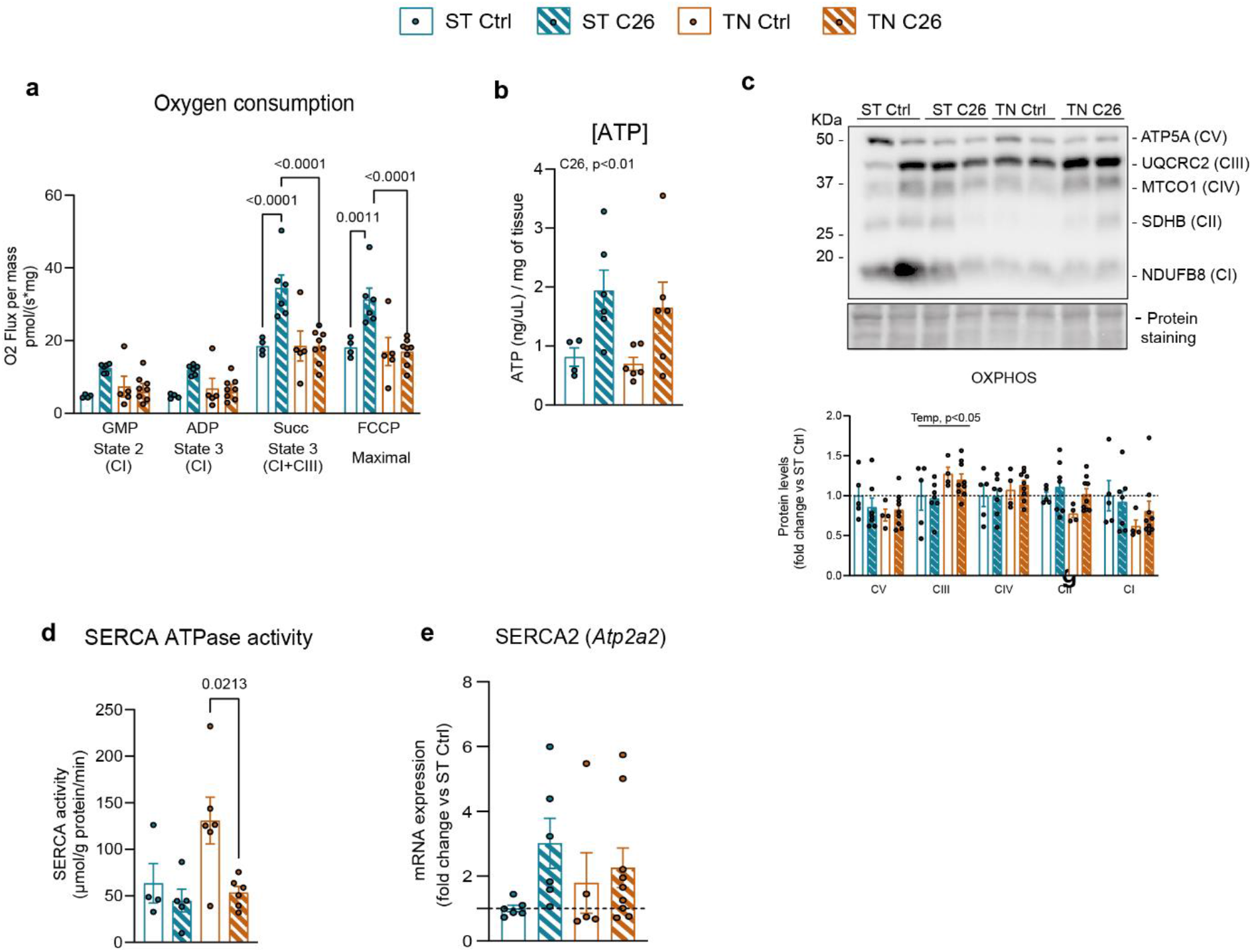
The cancer-induced molecular responses in BAT are strongly influenced by housing temperature. (**a**) Oxygen consumption in permeabilized BAT. (**b**) ATP concentration. (**c**) OXPHOS subunit representative immunoblots and quantification. (**d**) SERCA ATPase activity. (**e**) SERCA2 mRNA levels. Data are expressed as mean ± SE including individual values where applicable. Significance level was set at α=0.05 and only *p*<0.05 are shown. Main effects and interactions were calculated using two-way ANOVA and are shown only when Tukey’s post-hoc test analyses provided no statistical significance between experimental groups. (**a-e**) Two-way ANOVA test with Tukey’s post-hoc test. C26, main cancer effect; Temp, main temperature effect.

Preclinical mouse models have significantly advanced our understanding of cancer’s molecular and organismal impacts ^25^. However, we show that housing temperature crucially affects responses to murine cancer (Sup. Fig. 4d), highlighting the need to consider this factor in study design.

One of the most remarkable findings was that the bioenergetic adaptations to cancer occurring in BAT and SkM depended on housing temperature. Furthermore, BAT exhibited an enhanced sensitivity to housing temperature than SkM. We unprecedently show that BAT from C26 mice housed at ST, showing unaltered UCP1 protein content, presented remarkably increased O_2_consumption, recapitulating similar observations made in WAT of C26 mice housed at ST^9^. Yet, at TN, where UCP1 was downregulated, O_2_ consumption remained unchanged. Thus, these data indicate that the cancer-mediated adaptations in the mitochondrial respiratory capacity of BAT are sensitive to temperature, however, they do not correlate with UCP1 content and, consequently, to thermogenic activity. SkM showed the same, albeit less pronounced, trend in O_2_ consumption as BAT, yet contradicting reported preclinical studies^9^. This evidence suggests that the molecular mechanisms by which cancer results in temperature-independent SkM weakness might not involve the mild adaptations in SkM mitochondrial respiratory capacity occurring at different housing temperatures.

SkM weakness and induction of atrogenes preceded SkM mass loss in C26 mice and were not influenced by housing temperature. In contrast, fat mass was housing temperature-sensitive, as we observed a reduction in C26 mice only at TN, and the expression of genes involved in WAT or BAT function was differently regulated by cancer and temperature. Another key finding is that we observed decreased UCP1 gene expression in BAT and WAT of mice with moderate cachexia, recapitulating previous studies^15^. These findings feed the debate of whether CC is caused by increased UCP1 content in adipose tissue, leading to browning and exacerbated energy expenditure^10,16^.

We were also intrigued to find different temperature-sensitive regulations of SERCA ATPase activity, a process that is involved in thermogenic metabolism. Reduced SERCA ATPase activity is associated with age-related SkM atrophy and weakness^26^, and yet, despite all C26 mice exhibited SkM weakness, cancer induced an exacerbated SkM SERCA ATPase activity at ST, but not at TN. In contrast, BAT of C26 mice housed at ST exhibited a mild decrease in SERCA ATPase activity, and this response was enhanced in TN conditions. Thus, our data demonstrated that SkM and BAT SERCA ATPase activity is differently modulated depending on housing temperature upon cancer, which can lead to disparate consequences at the level of tissue metabolism.

Another set of key findings were that glucose metabolism was considerably affected by temperature in C26 mice, which could be due to relevant circulating factor playing a role in glycemic control, such as FGF21^27^. Interestingly, cancer increased plasma FGF21 levels, and this increase was enhanced upon TN. The same observation was made for the cytokine IL-6, which has also been shown to improve glucose metabolism in both healthy and obese mice^28^. Thus, it could be speculated that C26 mice housed at TN present improved glucose metabolism due to the synergistic and enhanced effect of FGF21 and IL-6. Yet, because of IL-6’s effects on SkM and adipose tissues, this also results in a detrimental impact further driving tissue dysfunction.

Altogether, our data demonstrate that housing temperature determines metabolic and molecular responses to cancer in mice, including alterations in body composition, glucose metabolism, and inflammatory responses. Our study also shows that housing temperature is a determinant factor in the cancer-induced cellular responses in mice, which are especially influenced in BAT. Due to known interactions between BAT and other tissues in both basal and during cancer^29^, we speculate that initial cancer-induced alterations in BAT bioenergetics could signal other tissues, especially SkM (Sup. Fig. 4d). Importantly, humans live in their TN zone, thereby not requiring thermogenesis from BAT^4^. Opposed to this notion, a recent study assessing both healthy subjects and patients with cancer, along with different tumor-bearing mouse models, demonstrated that a significant amount of BAT exists in humans, that it is activated upon cold exposure, and that cold-induced metabolic activity of BAT compromises tumor glucose uptake^30^. Our study, along with existing evidence, highlights the significant role of cold-induced BAT metabolism during cancer in both humans and mice, enhancing our understanding of translating preclinical findings to human pathophysiology.

## MATERIALS AND METHODS

### Animals

Fourteen-week old male BALB/C mice (Janvier) were maintained on a 12:12-h light-dark cycle with nesting material and *ad libitum* access to standard rodent chow diet (altromin no.1324; Chr. Pedersen, Denmark). Mice were randomly assigned to standard temperature (22 ºC ± 1 ºC) or thermoneutrality (30 ºC ± 1 ºC) in different rooms in the same animal facility. After a three-week acclimation period, mice were housed in pairs or triplets. Colon26 (C26) adenocarcinoma cells (Cytion, #400156) were cultured in RPMI 1640 medium (Gibco, #11875093, USA) supplemented with 10% fetal bovine serum (FBS, Sigma-Aldrich, #F0804, USA), 1% penicillin-streptomycin (ThermoFisher Scientific, #15140122, USA) (5% CO_2_, 37 ºC). Prior to inoculation into mice, C26 cells were trypsinized and washed twice with PBS. C26 cells were suspended in PBS at a final concentration of 5 x 10^6^ cells/mL. All mice were shaved the right flank one day prior to the inoculation. All mice were subcutaneously injected with PBS with or without 5 x 10^5^ C26 cells into the flank. Body weight and composition, as well as food intake, were monitored once a week during the first fourteen days after C26 cell inoculation, and twice a week after that. Mice developing ulcerations >5mm in diameter and tumors >14 mm in average of length and width (humane endpoint) were euthanized by cervical dislocation. Mice with tumors <0.5 g were excluded.

All experiments were approved by the Danish Animal Experimental Inspectorate (License; 2021-15-0201-01085).

### Body composition

Total, fat and lean body mass were measured weekly by nuclear magnetic resonance using an EchoMRI™ (USA). Body composition and weight were measured in all mice immediately prior to their dissection, and the final lean mass of C26 mice resulted from the subtraction of the tumor weight to the lean mass recorded prior dissection.

### Grip strength assessment

Grip strength was assessed two days prior the dissection. Mice were placed on the grip meter (Bioseb) with their forelimbs on the grid. Once on the apparatus, traction was applied via the tail. Each mouse had 5 runs, and the maximal recorded value of the resistance the mouse applied on the grid was chosen.

### *Ex vivo* force assessment

*Soleus* muscle was rapidly dissected, as previously described^31,32^, and incubated in heated (30°C) and oxygenated (95% O_2_, 5% CO_2_) Krebs-Ringer buffer (117 mM NaCl, 4.7 mM KCl, 2.5 mM CaCl_2_, 1.2 mM KH_2_PO_4_, 1.2 mM MgSO_4_, 24.6 mM NaHCO_3_, 0.1% bovine serum albumin, and 5 mM glucose) in a myograph system (820MS DT, Denmark). Prior to the experiment, optimal muscle length was determined by finding the length of maximal isometric twitch response (pulse voltage 14 V, pulse width 0.5 ms). Single twitch assessment was conducted (three stimuli of 14 V and 0.5 ms pulse width separated by >30 s and then averaged) to determined maximal twitch force, time to peak force development, and half relaxation time. For tetanic contractions (pulse voltage 14 V, pulse width 0.2 ms, interval 6.50 ms, frequency 150 Hz, 75 pulse counts in each train, 500 ms total train time, 5,000 ms pause between trains for a total of 110 trains), the area under the curve was measured to assess total force production by summating the products of peak force and contraction duration from each tetanic contraction. Absolute force measurements were normalized to muscle cross sectional area to find the specific force (muscle mass divided by the product of the muscle density coefficient (1.06 g/cm^3^), optimal muscle length, and the muscle-specific fiber length coefficient (0.71 for *soleus*^33^).

### Glucose tolerance test

Glucose (Sigma-Aldrich, #1083421000) was dissolved in saline at a concentration of 0.2 g/mL and intraperitoneally injected into 4-h-fasted mice (fasting single-housed from 7:00 am) at a dose 2 g/kg body weight. At time points 0, 20, 40, 60 and 90 min, blood glucose was measured using a glucometer (Bayer Contour; Bayer, Münchenbuchsee, Switzerland). At time points 0 and 20, blood was collected from the tail using EDTA-containing collection tubes (Microvette, VWR).

### Plasma lipidomics

Quantification of lipid species in plasma samples was performed as previously described by Shashikadze et al. 2023^34^. Briefly, after addition of 25 µL of a mix of 77 deuterated internal standards, plasma samples were extracted twice using water, methanol (MeOH) and methyl tert-butyl ether (MTBE)^35^. The combined organic phases were evaporated to dryness with N_2_ and reconstituted in 275 µL running solvent (10 mM ammonium acetate in Dichloromethane:MeOH (50:50, v/v)).75 µL were injected into a system consisting of a SCIEX Exion UHPLC-systemcoupled to a SCIEX QTRAP 6500+ mass spectrometer equipped with a SelexION differential ion mobility interface (SCIEX, Darmstadt, Germany) operated with Analyst 1.6.3. Details about the MRM-based quantification method can be found in Shashikadze et al. 2023^34^. The shotgun lipidomics raw data set contained 989 individual lipid species. Data were subsequently pre-processed using R (version 4.2.1). A multi-step procedure was applied to ensure high data quality: in the first step of this quality control (QC) procedure, lipids with missing values in more than 35% in the pool samples were discarded from the data set (n = 114). In the second step, the group-specific missingness was evaluated i.e., whether a specific lipid is observed in only one of the biological groups. Lipids exhibiting a groupwise missingness of 50% in all groups were discarded from the data set (n = 1). Next, lipids with a coefficient of variation >25%, determined by the QC-pool samples, were removed from the data set (n = 32). In a final QC step, lipids having a dispersion ratio^36^. 50% were removed (n = 65). After quality control, 777 lipid species remained in the data set, which contained 319 missing values (equivalent to 1.4% of the data set). Missing values were imputed using the k-nearest-neighbor obs-sel approach with k = 10 nearest-neighbors^37^.

### Plasma measurements

Plasma levels of circulating cytokines and hormones were measured using a customized multiplex assay kit by V-PLEX platform (Mesoscale, Maryland, USA) and following the manufacturer’s instructions. In this kit, we included the cytokines IL-6, TNF-α, IFN-γ and IL-1β, and the hormones FGF21, insulin and leptin. Plasma GDF-15 was measured using the Mouse/Rat GDF-15 Quantikine ELISA Kit (R&D Systems, MGD150).

### Respirometry

BAT tissues and *quadriceps* muscles were placed in ice-cold isolation solution BIOPS (OROBOROS Instruments, Innsbruck, Austria) right upon dissection. BAT tissue was further dissected into small pieces using scissors while muscle fibers were gently mechanically separated using needles. Then tissues were permeabilized in either 2.5 μg/mL of digitonin or 50 μg/mL of saponin for BAT or muscle fibers, respectively, for 30 min on ice and on an orbital shaker. This was followed by 2 washes (10 min) in mitochondrial respiration medium MIRO5 (OROBOROS Instruments, Innsbruck, Austria). Then ∼2-3 mg (wet weight) of muscle fibers or BAT tissue were transferred to the oxygraph-2k (O2k; OROBOROS Instruments, Innsbruck, Austria) containing respiration medium. All respiration measurements were performed at 37 ºC and in hyperoxia conditions ([O_2_: 200-450 umol/L). We assessed the following protocol: addition of 10 mM glutamate, 2 mM malate and 5 mM sodium pyruvate to assess resting respiration (state II, absence of adenylates), followed by addition of 2.5 mM ADP to assess state III. Then we examined uncoupling control by adding several steps of 1 μM of the uncoupler [(3-chlorophenyl)hydrozono] malononitrile (CCCP) until optimum concentration. Inhibition of complexes I and III was achieved by adding 0.5 μM of rotenone, followed by 2.5 μM of antimycin A to evaluate non-mitochondrial respiration.

### Tissue processing

All tissues were quickly dissected and snap frozen in liquid nitrogen. BAT and skeletal muscles were pulverized using a Bessman type tissue pulverizer (Cellcrusher kit) on dry ice. For RNA extraction, tissues were homogenized in QIAzol lysis reagent (#79306 Qiagen) and three ceramic beads. For ATP determination, tissues were homogenized in 150 μL of ATP Assay Buffer from the ATP Colorimetric/Fluorimetric Assay Kit (Sigma-Aldrich, MAK190). For protein extraction, tissues were homogenized in ice cold homogenization buffer (10% glycerol, 1% NP-40, 20 mM sodium pyrophosphate, 150 mM NaCl, 50 mM HEPES (pH 7.5), 20 mM β-glycerophosphate, 10 mM NaF, 2mM phenylmethylsulfonyl fluoride (PMSF), 1 mM EDTA (pH 8.0), 2 mM Na_3_VO_4_, 10 μg/mL leupeptin, 10 μg/mL aproptinin, 3 mM benzamidine) with a steel bead^7,17^ using the TissueLyser II bead mill (Qiagen, USA) for 1 min at 30 Hz.

### Gene expression analyses

RNA extraction was performed using the RNeasy Mini Kit (Qiagen #74106) following the manufacturer’s instructions. RNA purity was measured with the Nanodrop™ 2000/20000c spectrometer and the ND1000 software (ThermoFisher Scientific). Reverse transcription reaction of was performed using 4 μL of qScript Ultra SuperMix (#95217, Quantabio) per 500 – 1000 ng of RNA. Gene expression was quantified based on real-time quantitative PCR using SYBR green (Applied Biosystems) and the QuantStudio 6 and 7 Real-Time PCR System (Applied Biosystems), using the default protocol for comparative Ct studies. All measurements were normalized to housekeeping genes, namely *b-actin* or *36b4*. All primers used and their sources are listed in Supplementary Table 1.

### [ATP] determination

Prior to the ATP determination assay, the deproteinizing sample preparation kit – tricarboxylic acid (TCA) (Abcam, ab204708) was used in pulverized BAT and *triceps* muscles to remove macromolecules that interfere with further ATP quantification. Homogenized tissues were incubated with 15 μL of ice-cold TCA in ice for 15 min, and then centrifuged at 12000 x *g* for 5 minutes at 4 ºC. Supernatants were collected and excess of TCA was neutralized by adding 10 μL of cold neutralization solution. Samples were then allowed to vent as CO_2_ might be formed on ice for 5 min and stored at −80 ºC until further ATP determination. The ATP Assay Kit (Sigma-Aldrich, #MAK190) was performed by measuring fluorescence following the manufacturer’s instructions.

### Immunoblotting

Upon tissue homogenization for protein extraction, samples were subjected to end-over-end rotation for 30 min at 4 ºC, and then centrifuged at 9500 x *g* for 20 min at 4 ºC. Supernatants (protein extracts) were collected and stored at −80 ºC. The latter two steps (centrifugation and lysate collection) were performed three times in BAT to avoid contamination of fatty acids. Protein concentration was measured using the bichinchoninic acid (BCA, ThermoFischer #23225) method with bovine serum albumin (BSA, Sigma-Aldrich #P0834) as standard. 20 ug of protein extracts were resolved in 7 %, 10 % or 12.5 % Mini-PROTEAN precast acylamide gels (BioRad 4561026, 4561036, 4561046) for SDS-PAGE and transferred to polyvinylidene difluoride membranes (Immobilon Transfer Membrane, Millipore) and then blocked in Tris-buffered saline (TBS)-Tween 20 containing 2% milk protein for 30 min at room temperature. Membranes were incubated with primary antibodies overnight at 4 ºC, followed by incubation with horseradish peroxidase-conjugated secondary antibody for 60 min at room temperature. Pierce™ Reversible Stain (ThermoFisher #24585) or a Coomassie Brilliant Blue R-250 staining (Bio-Rad, #1610400) was used as loading control. Bands were visualized using the Bio-Rad ChemiDoc MP Imaging System and enhanced chemiluminescence (ECL+, Amersham Biosciences). Band densitometry was carried out using Quantity One (Bio-Rad). Primary antibodies used are specified in Supplementary table 2.

### SERCA ATPase activity assay

To determine SERCA ATPase activity, an enzyme-linked spectrophotometric assay was used, as previously described^38–40^, in the absence of ionophore (i.e., presence of Ca^2+^ gradient). Briefly, BAT or *triceps* muscle homogenates were added to a reaction buffer with maximally activating Ca^2+^ concentrations (*p*Ca 5) and plated in duplicate in a 96-well plate. The reaction was initiated with the addition of NADH. The rate of NADH disappearance, which occurs in a 1:1 ratio with ATP hydrolysis, was measured over 30 min. A baseline rate was obtained using the SERCA specific inhibitor, cyclopiazonic acid, which was subtracted from all other rates to obtain SERCA-specific ATPase activity. SERCA activity was then normalized to sample protein concentration.

### Statistics and graphics

The data presented here were collected using Excel 2016 and were analyzed using GraphPad Prism version 10.1.1 for Windows (GraphPad Software, San Diego, California USA, www.graphpad.com). Data are expressed as mean ± SE including individual values where applicable. Significance level was set at α=0.05 and only *p*<0.05 are shown. Main effects and interactions between two conditions (temperature and cancer) were calculated using Two-way ANOVA and Tukey’s post-hoc tests were only calculated and shown when main effects/interactions were significant. Data plots were generated using GraphPad Prism version 10.1.1 for Windows (GraphPad Software, San Diego, California USA, www.graphpad.com). The graphics shown in Sup. Fig. 4d were generated with ©BioRender – biorender.com (Toronto, Canada)

## ACKNOWLEDGMENTS

We acknowledge the contribution of B. Bolmgren and E.A. Richter (Department of Nutrition, Exercise and Sports, Faculty of Science, University of Copenhagen, Denmark) in analyzing GDF-15 levels in plasma samples.

## Author contributions

Conceptualization: A.I and L.S. Methodology: A.I., T.C.P.P., J.B., A.M.E., M.H., C.H.F.H., R.M-V, S.L., and L.S. Investigation: A.I., E.F., T.C.P.P., J.B., A.M.E., M.H., F.R., C.H.F.H., A-S.R.J., N.R.A., L-H-C., R.M-V., M.S.A., S.H.R., J.L.M., S.G., S.L., Z.G-H., T.E.J., M.R., J.T.T., V.A.F. and L.S. Visualization: A.I. and L.S. Supervision: L.S. Writing (original draft): A.I. and L.S. Writing (review and editing): A.I., E.F., T.C.P.P., J.B., A.M.E., M.H., F.R., C.H.F.H., A-S.R.J., N.R.A., L-H-C., R.M-V., M.S.A., S.H.R., J.L.M., S.G., S.L., Z.G-H., T.E.J., M.R., J.T.T., V.A.F., and L.S.

