## Supplementary information for "Housing temperature dictates the systemic and tissue-specific molecular responses to cancer in mice"

### Figures

#### Supplementary Figure 1

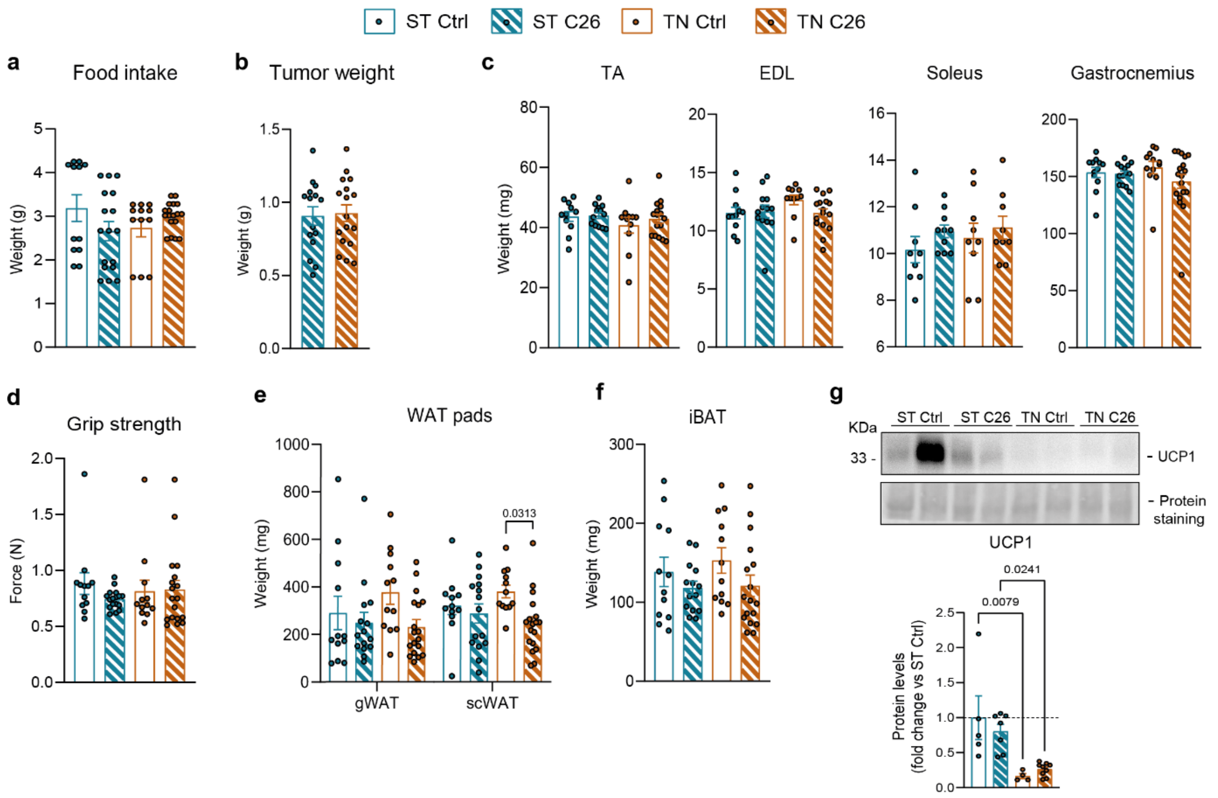


**Supplementary Figure 1.** (**a**) Food intake quantification. (**b**) Tumor weight. (**c**) Muscle weights. (**d**) Grip strength quantification. (**e**) Weights of WAT depots. (**f**) iBAT weights. (**g**) Representative immunoblot of UCP1 and band quantification in BAT. Data are expressed as mean ± SE including individual values where applicable. Significance level was set at α=0.05 and only $p<0.05$ are shown. Main effects and interactions were calculated using two-way ANOVA and are shown only when Tukey's post-hoc test analyses provided no statistical significance between experimental groups. (**a, c-g**) Two-way ANOVA with Tukey's post-hoc test. (**b**) Student's T test.

### Supplementary Figure 2

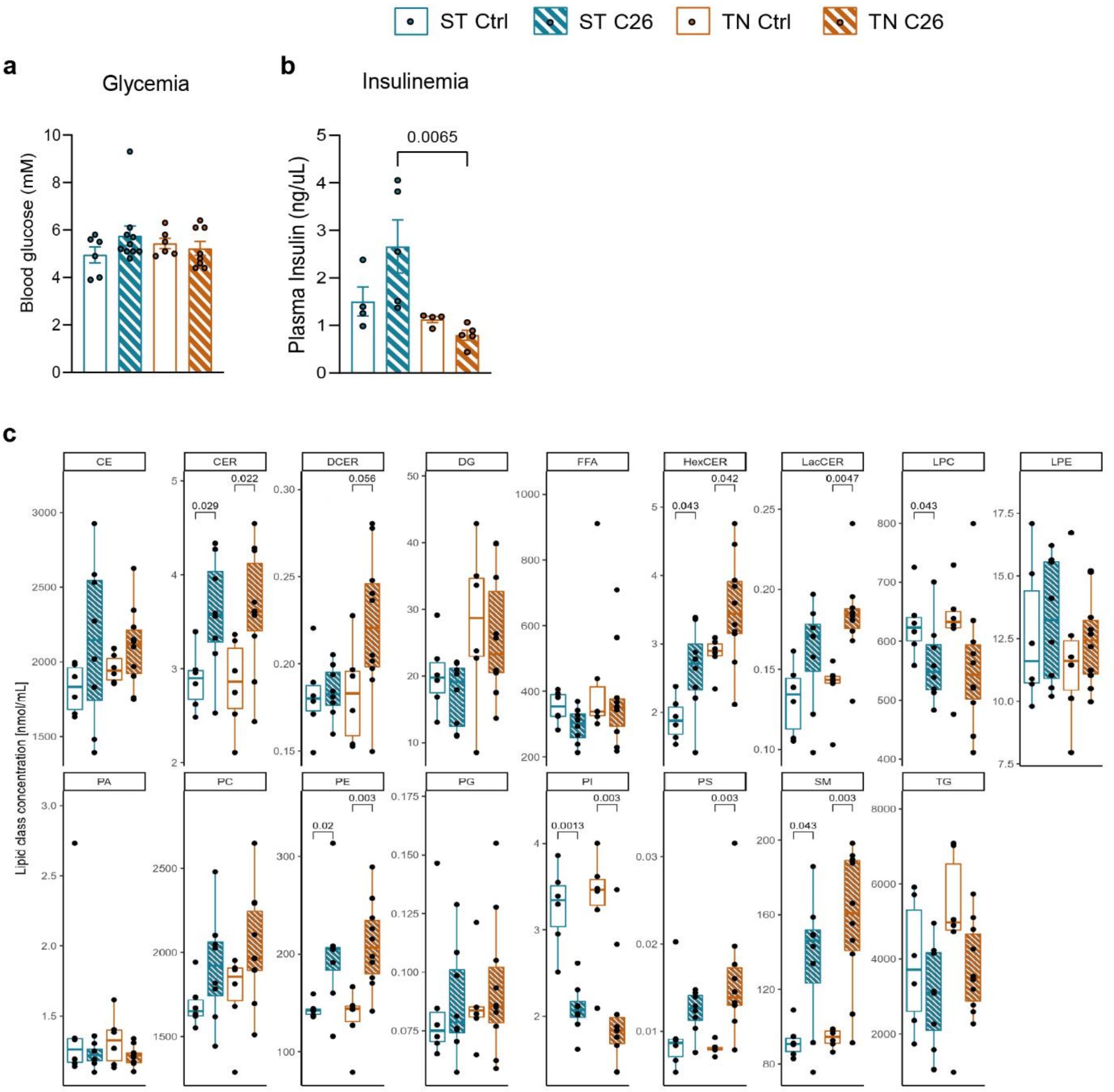


**Supplementary Figure 2.** (**a**) Fasting blood glucose. (**b**) Fasting blood insulin. (**c**) Plasma lipid class sum concentration. Data are expressed as mean ± SE including individual values where applicable. Significance level was set at α=0.05 and only $p$<0.05 are shown. Main effects and interactions were calculated using two-way ANOVA and are shown only when Tukey's post-hoc test analyses provided no statistical significance between experimental groups. (**a, b**) Two-way ANOVA test with Tukey's post-hoc test. (**c**) Wilcoxon test.

**Supplementary Figure 3**

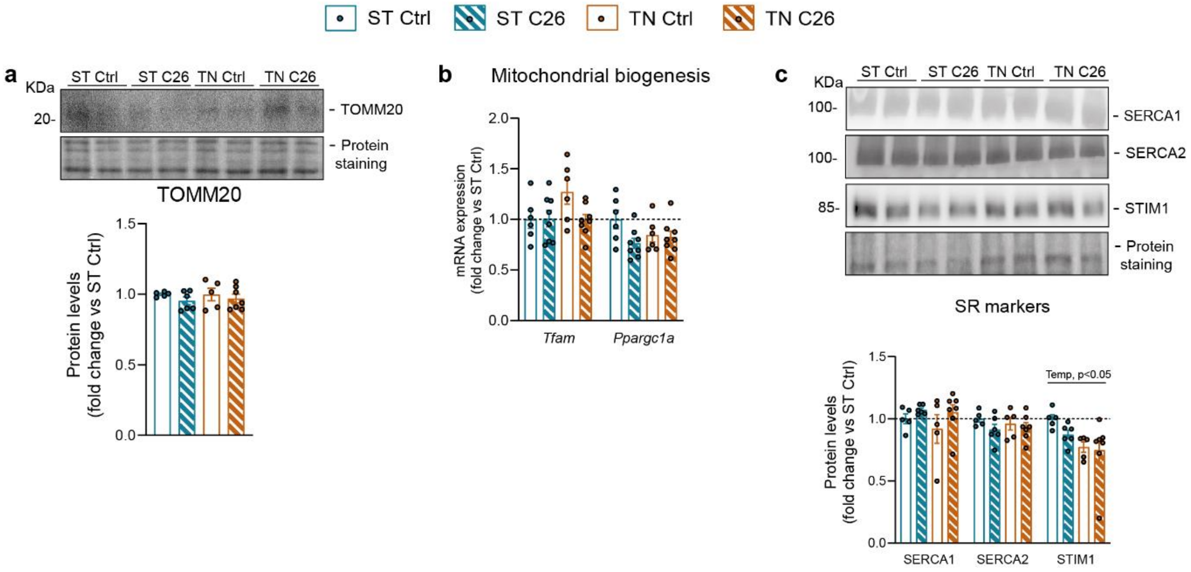


**Supplementary Figure 3.** (**a**) Representative immunoblot of TOMM20 and band quantification in SkM. (**b**) *Tfam* and *Ppargc1a* mRNA levels SkM. (**c**) Representative immunoblots of ER/SR markers and band quantification in SkM. Data are expressed as mean ± SE including individual values where applicable. Significance level was set at α=0.05 and only $p<0.05$ are shown. Main effects and interactions were calculated using two-way ANOVA and are shown only when Tukey's post-hoc test analyses provided no statistical significance between experimental groups. (**a-c**) Two-way ANOVA test with Tukey's post-hoc test. Temp, main temperature effect.

**Supplementary Figure 4**

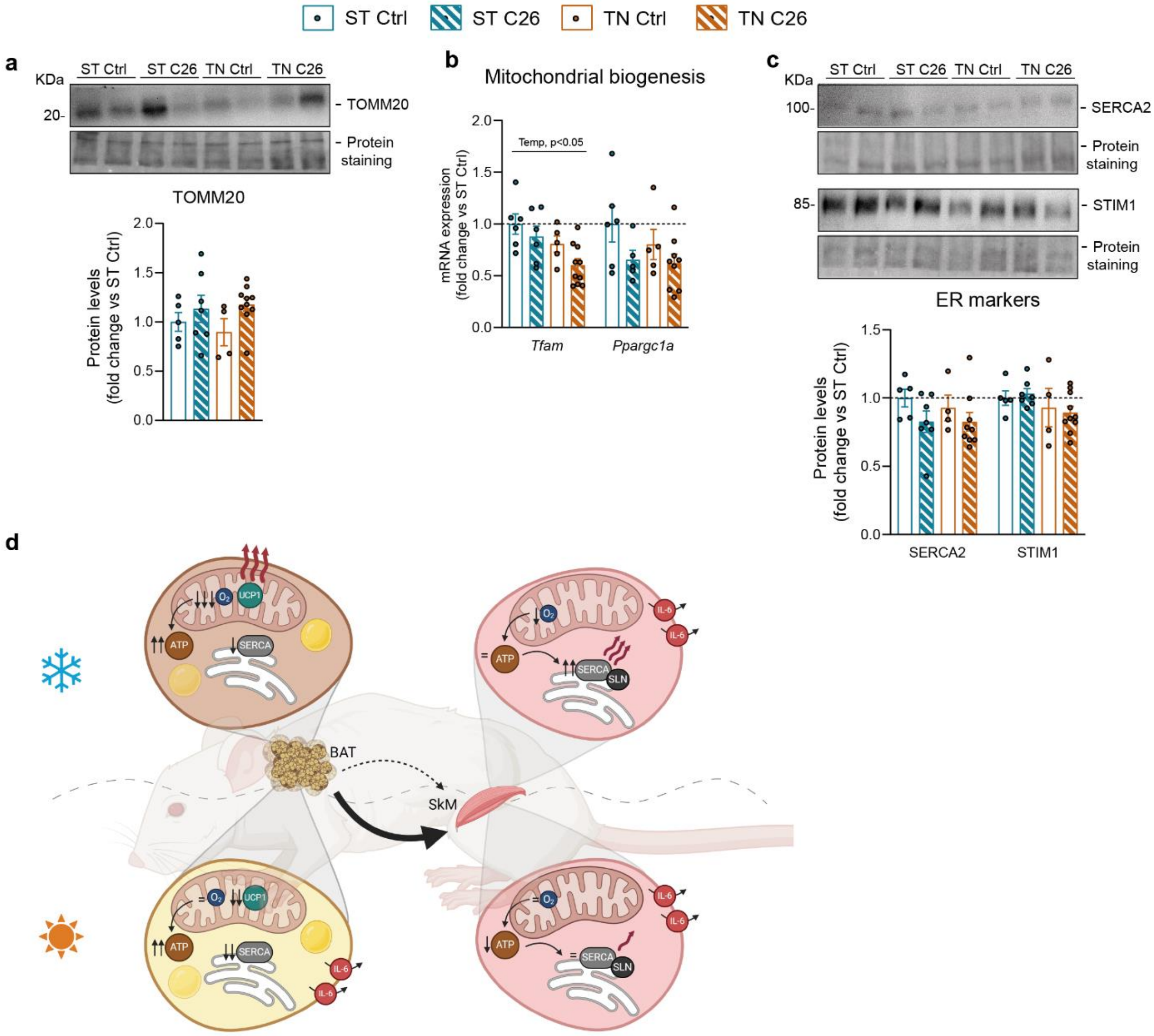


**Supplementary Figure 4.** (**b**) Representative immunoblot of TOMM20 and band quantification in BAT. (**b**) *Tfam* and *Ppargc1a* mRNA levels BAT. (**c**) Representative immunoblots of ER markers and band quantification in BAT. (**d**) Graphical abstract of the current working model based on the results obtained in this study. Data are expressed as mean ± SE including individual values where applicable. Significance level was set at α=0.05 and only $p$<0.05 are shown. Main effects and interactions were calculated using two-way ANOVA and are shown only when Tukey's post-hoc test analyses provided no statistical significance between experimental groups. (**a-c**) Two-way ANOVA test with Tukey's post-hoc test. Temp, main temperature effect.

**Supplementary Table 1: SYBR green mouse primers**

| *Targeted gene* | *Forward sequence* | *Reverse sequence* | *Source* |
|---|---|---|---|
| *4ebp1* | CACGCTCTTCAGCACCAC | GGAGGCTCATCGCTGGTAG | Ninfali et al 2018 doi: 10.1093/nar/gky835 |
| *Fbxo32* | GCAAACACTGCCACATTCTCTC | CTTGAGGGGAAAGTGAGACG | Ninfali et al 2018 doi: 10.1093/nar/gky835 |
| *Trim63* | TGTCTGGAGGTCGTTTCCG | ATGCCGGTCCATGATCACTT | Ninfali et al 2018 doi: 10.1093/nar/gky835 |
| *Slc25a7* | AGGCTTCCAGTACCATTAGGT | CTGAGTGAGGCAAAGCTGATTT | Rupar et al 2023 doi: 10.1111/febs.16716 |
| *Cebpa* | CAAGAACAGCAACGAGTACCG | GTCACTGGTCAACTCCAGCAC | Primerbank |
| *Cepbd* | CGACTTCAGCGCCTACATTGA | CTAGCGACAGACCCCACAC | Primerbank |
| *Leptin* | GAGACCCCTGTGTCGGTTC | CTGCGTGTGTGAAATGTCATTG | Primerbank |
| *Dio2* | AGTCAAGAAGGTGGCATTCG | ACAGCTTCCTCCTAGATGCCT | Yoshizawa et al 2022 doi: 10.1038/s41467-022-35219-z |
| *Elvol3* | TTGGGGATAGGGGGTGTGTG | TCTCCCCTCCCCTCCAAGTC | Yoshizawa et al 2022 doi: 10.1038/s41467-022-35219-z |
| *Sirt7* | TGCCAGGCACTTGGTTGTCT | TAGGCTCCGCTTCGCTTAGG | Yoshizawa et al 2022 doi: 10.1038/s41467-022-35219-z |
| *Il6* | TAGTCCTTCCTACCCCAATTTCC | TTGGTCCTTAGCCACTCCTTC | Irazoki et al 2023 doi: 10.1038/s41467-022-35732-1 |
| *Tfam* | ATTCCGAAGTGTTTTTCCAGCA | TCTGAAAGTTTTGCATCTGGGT | Primerbank |
| *Ppargc1a* | TATGGAGTGACATAGAGTGTGCT | CCACTTCAATCCACCCAGAAAG | Primerbank |
| *Atp2a1* | TGTTTGTCCTATTTCGGGGTG | AATCCGCACAAGCAGGTCTTC | Primerbank |
| *Atp2a2* | GAGAACGCTCACACAAAGACC | CAATTCGTTGGAGCCCCAT | Primerbank |
| *Sln* | GCTCCTCTTCAGGAAGTGAAG | TGGCCCCTCAGTATTGGTAGG | Pant et al 2015 doi: 10.1242/jeb.119164 |
| *b-actin* | GGTCATCACTATTGGCAACGA | GTCAGCAATGCCTGGGTACA | Irazoki et al 2023 doi: 10.1038/s41467-022-35732-1 |
| *36b4* | TCATCCAGCAGGTGTTTGACA | GGCACCGAGGCAACAGTT | Self-designed |

**Supplementary Table 2: Primary antibodies**

| *Target* | *Catalog no.* | *Company* | *Dilution* |
|---|---|---|---|
| Anti-OXPHOS cocktail | MS604-300 | Abcam | 1:1000 |
| Anti-UCP1 | Ab155117 | Abcam | 1:1000 |
| Anti-TOMM20 | Sc-17764 | Santa Cruz Biotechnology | 1:1000 |
| Anti-SERCA1 | MA3-912 | Invitrogen | 1:1000 |
| Anti-SERCA2 | MA3-910 | Invitrogen | 1:1000 |
| Anti-STIM1 | 4916 | Cell Signaling | 1:1000 |
